# Maternal investment evolves with larger body size and higher diversification rate in sharks and rays

**DOI:** 10.1101/2022.01.05.475057

**Authors:** Christopher G Mull, Matthew W Pennell, Kara E Yopak, Nicholas K Dulvy

## Abstract

Across vertebrates, live-bearing has evolved at least 150 times from ancestral egg-laying into a diverse array of forms and degrees of prepartum maternal investment.^1,2^ A key question is how this reproductive diversity arose and whether reproductive diversification underlies species diversification?^3–11^ To test these questions, we evaluate the most basal jawed vertebrates, sharks, rays, chimaeras of Class Chondrichthyans, which have one of the greatest ranges of reproductive and ecological diversity among vertebrates.^2,12^ We reconstructed the sequence of reproductive mode evolution across a time-calibrated molecular phylogeny of 610 chondrichthyans.^13^ We find egg-laying is ancestral, and live-bearing evolved at least seven times. Matrotrophy (i.e. maternal contributions beyond the yolk) evolved at least 15 times, with evidence of one reversal. In sharks, transitions to live-bearing and matrotrophy are more prevalent in larger-bodied species in the tropics. Further, the evolution of live-bearing is associated with a near-doubling of the diversification rate, but, there is only a small increase in diversification associated with the appearance of matrotrophy and increased rates of speciation are associated with the colonization of novel habitats, contrary to what has been demonstrated in teleosts.^3,4^ This highlights a potential key difference between chondrichthyans and other fishes, specifically a slower rate of reproductive isolation following speciation, suggesting different rate-limiting mechanisms for diversification between these clades.^14^ The chondrichthyan diversification and radiation, particularly throughout the shallow tropical shelf seas and oceanic pelagic habitats, appears to be associated with the evolution of live-bearing and the proliferation of a wide range of maternal investment in developing offspring.

## Results and Discussion

We reveal the first chondrichthyan was an egg-layer and there have been numerous transitions toward live-bearing and matrotrophy (Figure 1). The evolution of live-bearing and matrotrophy covaries with increasing body size and is more prevalent in shallow waters of tropical latitudes (Figure 2). Further, the evolution of live-bearing, and to a lesser extent matrotrophy, appears to have resulted in greater species diversification (Figure 3,4). Next we unpack these findings by considering three questions: What is the sequence of reproductive mode evolution? What ecological factors have driven the evolution of live-bearing and matrotrophy? Is chondrichthyan speciation and diversification explained by viviparity-driven conflict or by novel ecological opportunity?

**Figure 1.**
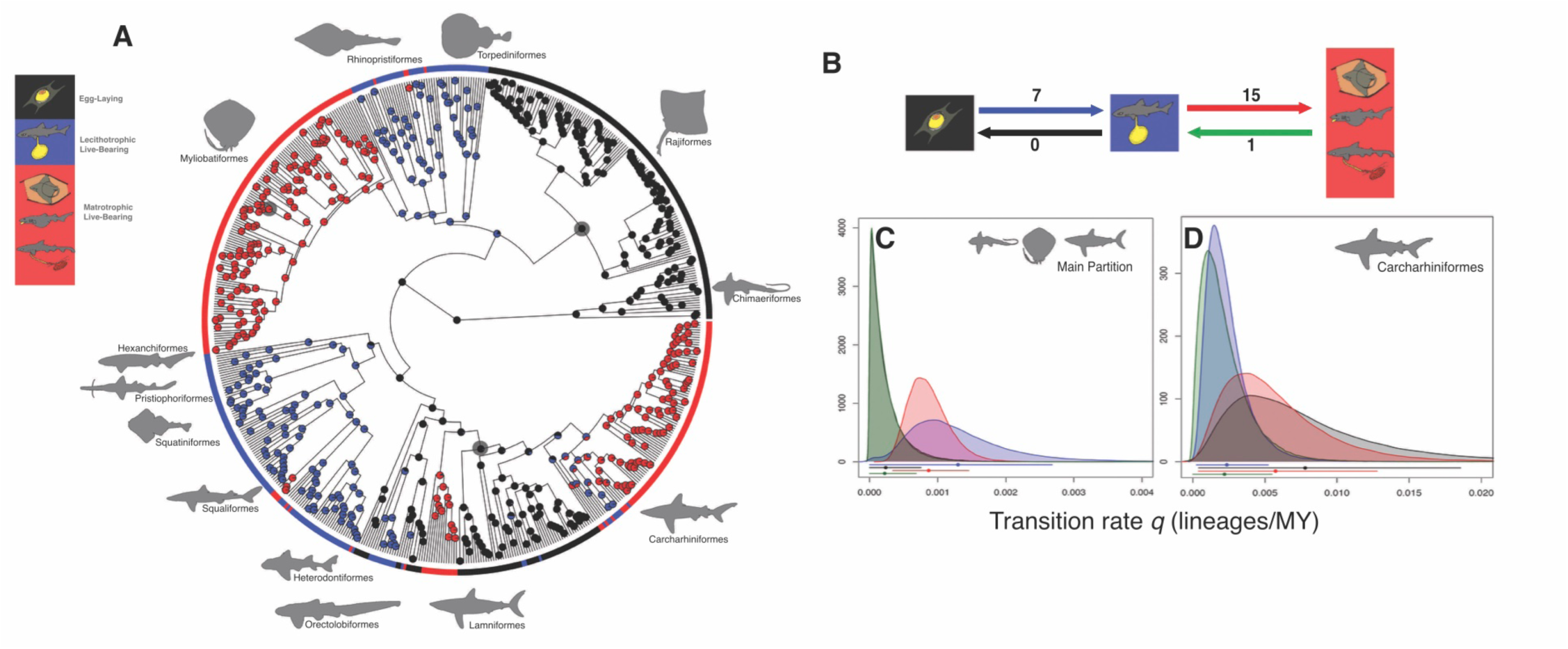
(A) Ancestral state reconstruction reproductive mode on a representative tree of 610 species of chondrichthyans. Pie symbols represent the likelihood of the character state for each node being egg-laying (black), live-bearing (blue), or matrotrophic (red). Dark grey symbols denote the partitions encompassing diversification rate shifts. Silhouettes depict representative species from the major orders. (B) The number evolutionary transitions in reproductive mode across chondrichthyans and transition rates between modes in (C) main partition of the tree and (D) within Carcharhiniformes. Origins of live-bearing from egg-laying are depicted in blue with reversals in black, and origins of matrotrophy from lecithotrophic live-bearing are depicted in red with reversals in green. Bars and shaded regions in represent the 95% posterior density of transition rate estimates.

**Figure 2.**
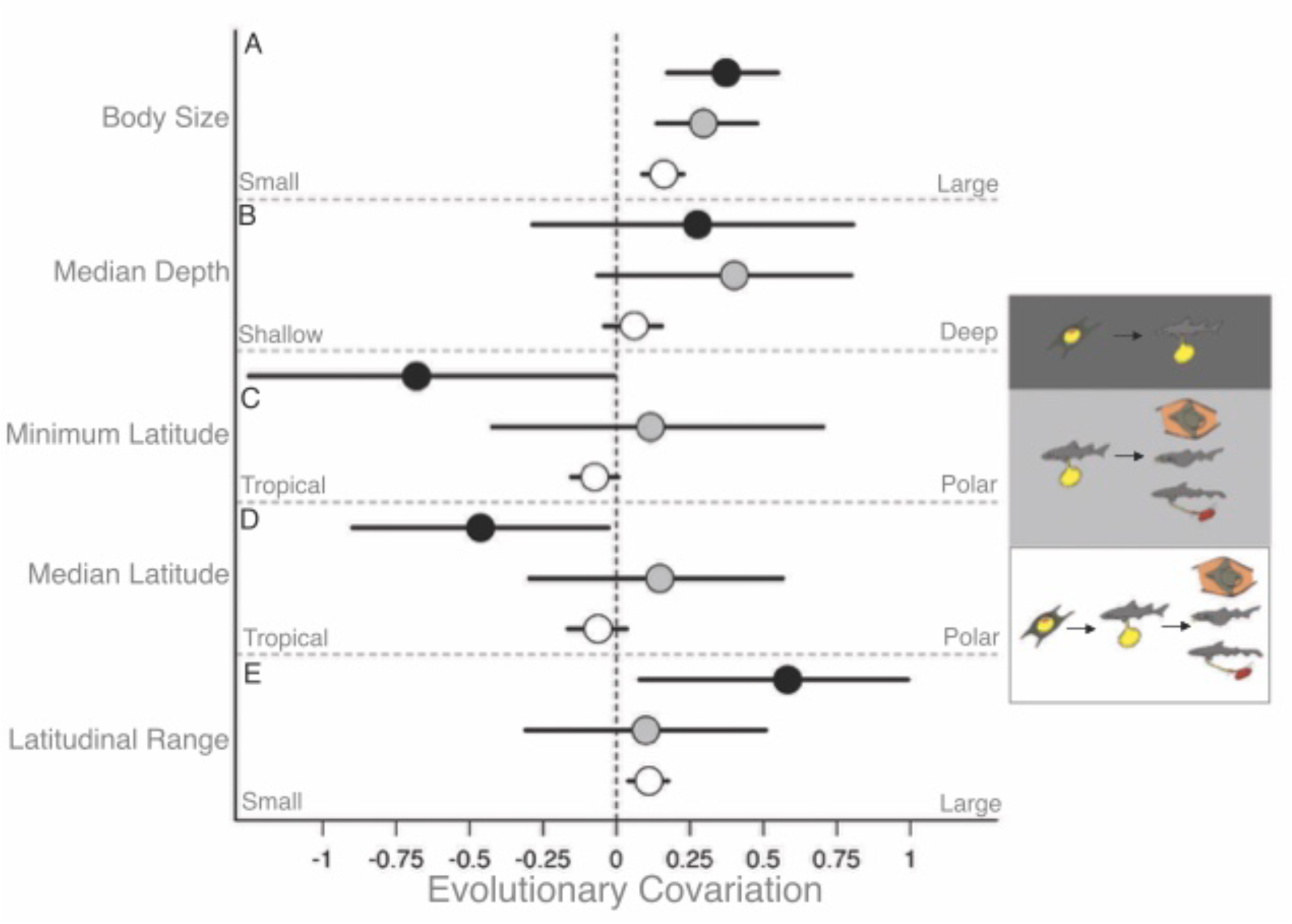
Coefficient plots of evolutionary covariation between (A) body size, (b) median depth, (c) minimum latitude, (d) median latitude, and (e) latitudinal range from MCMCglmm models. Black circles denotes egg-laying vs. live-bearing, grey circles denote live-bearing vs matrotrophy, and open circles denote reproductive mode as an ordinal variable with all three character states. Horizontal bars represent the 95% confidence intervals of the mean posterior estimate.

**Figure 3.**
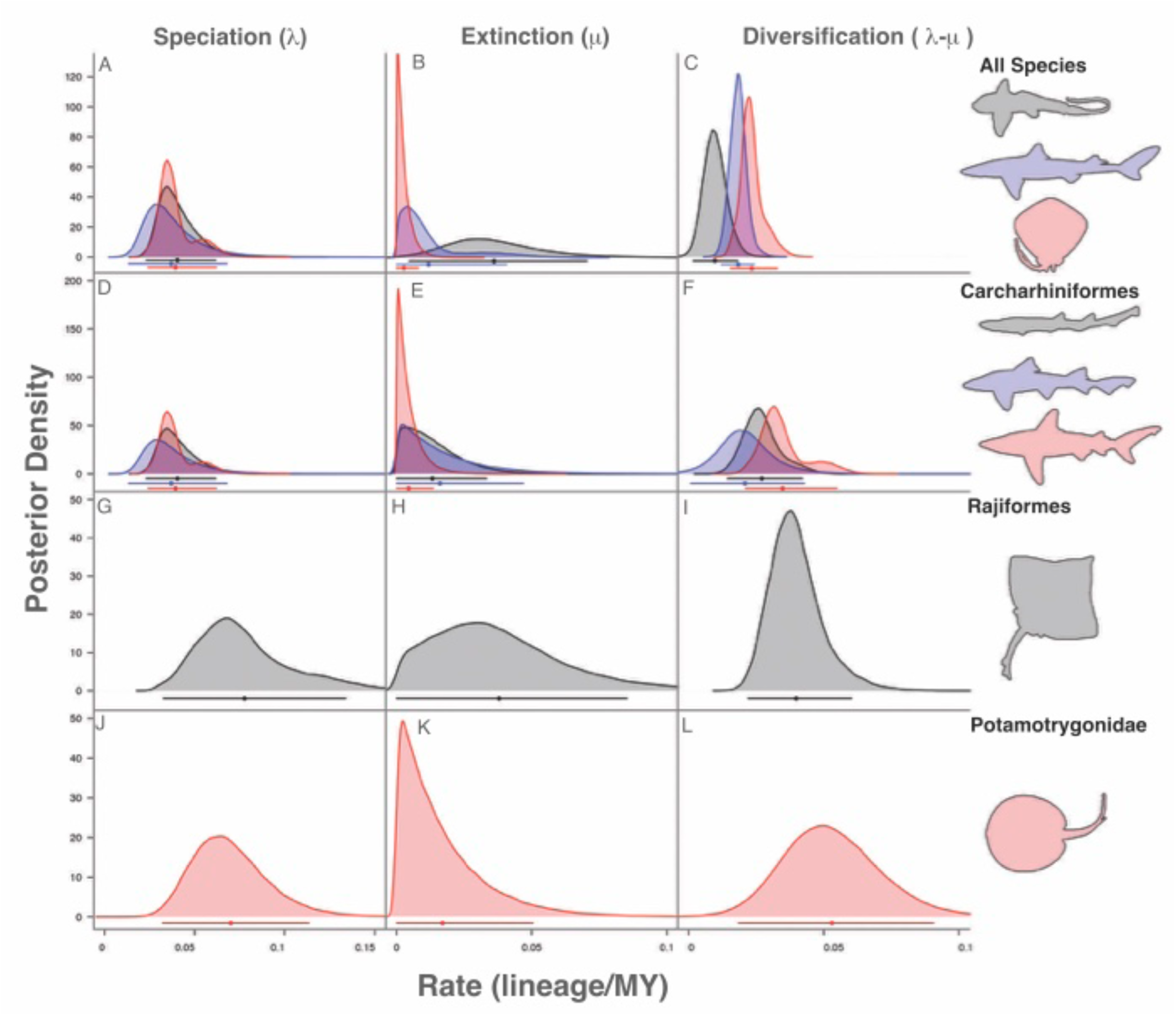
Posterior densities of parameter estimates from MuSSE model showing state dependent speciation (λ), extinction (μ), and net diversification rate (*r*) from the main partition of the tree (A-C), within the order Carcharhiniformes (D-F), within the order Rajiformes (G-I), and within the family Potamotrygonidae (J-L). Egg-laying is depicted in black, live-bearing in blue, and matrotrophy in red. Bars and circles represent the mean and 95% confidence interval of the posterior mean estimate.

**Figure 4.**
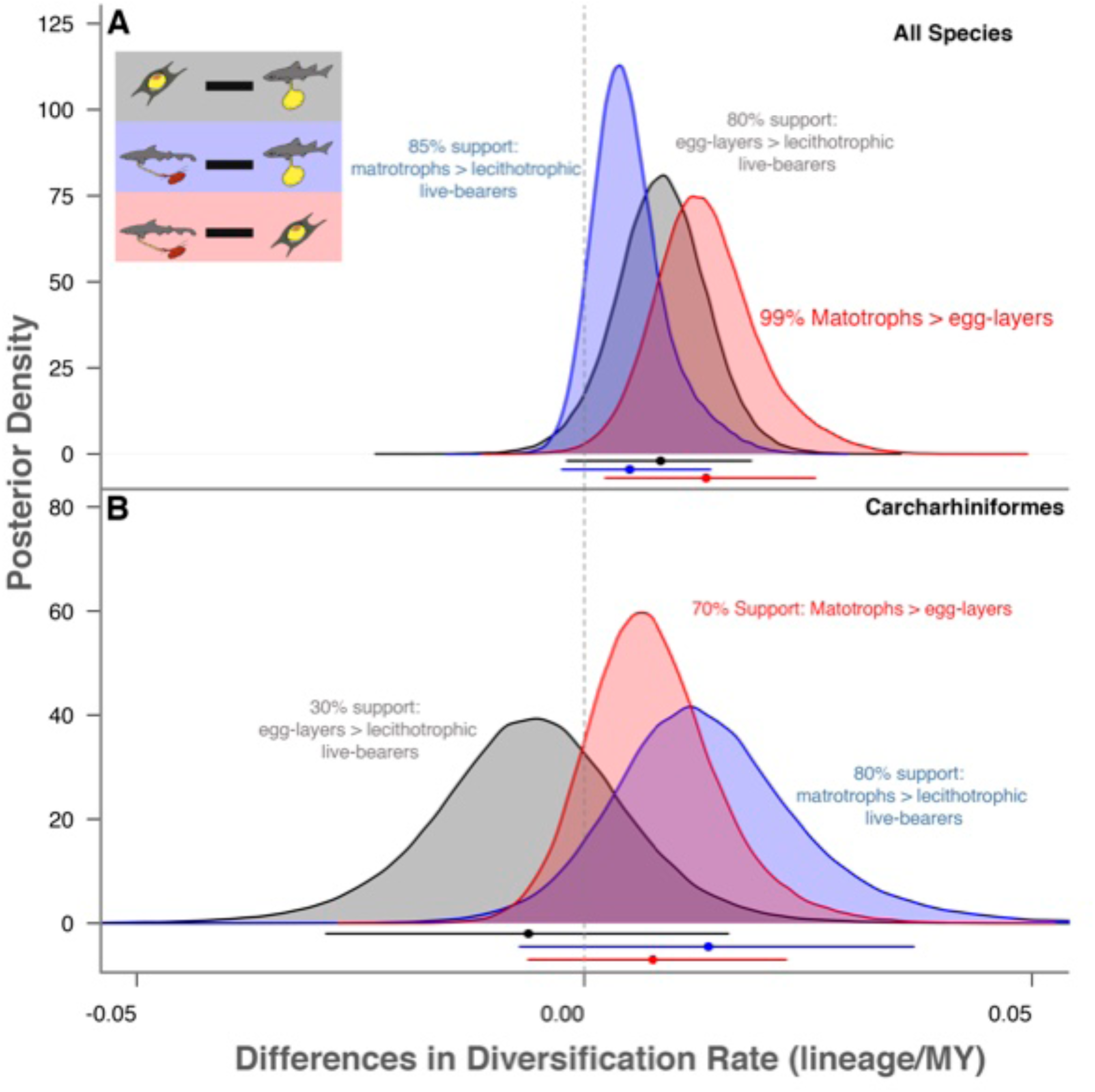
Differences between posterior densities of parameter estimates from MuSSE model showing the differences in reproductive mode dependent net diversification rates (*r*) from the main partition of the tree (A), and within the order Carcharhiniformes (B). Live-bearing relative to egg-laying is depicted in black, matrotrophy relative to live-bearing in blue, and matrotrophy relative to egg-laying in red. Bars and circles represent the mean and 95% confidence interval of the posterior mean estimate. The vertical lines denote no difference between the diversification rates for the reproductive modes being compared.

### The sequence of reproductive mode evolution

The first chondrichthyans almost certainly laid eggs, as there is a high level of support for egg-laying (oviparity) as the ancestral state of reproductive mode in chondrichthyans (>99% probability; Figure 1A; see Supplementary Table 1 for a full description of each reproductive mode). There are multiple independent origins of live-bearing (seven) and matrotrophy (15), at a range of taxonomic levels from the superordinal to subgeneric level, with few instances of reversals (one reversal from matrotrophy to lecithotrophic live-bearing; Figure 1B-D). Specifically, live-bearing appears to have evolved from egg-laying at: (a) base of Rhinopristiformes and Myliobatiformes, (b) base of Squalomorphii, (c) base of clade encompassing Brachaeluridae, Orectolobidae, and Rhincodontidae, (d) base of Ginglymostomatidae, (e) within the genus *Bythaelurus*, (f) in *Galeus polli*, and (g) basal to clade encompassing Pseudotriakidae, Triakidae, Hemigaleidae, and Carcharhinidae (Figure 1A). We found no evidence of reversals from live-bearing to egg-laying (Figure 1B). Previous analyses of reproductive evolution in chondrichthyans came to this conclusion based on the available morphological phylogenetic hypotheses, that incorrectly inferred that rays (and skates) as a highly derived group within the Squalimorph sharks^15^, but the emerging consensus from molecular data is that rays and sharks are sister taxa.^13,16^ With this new molecular tree, we find transitions in reproductive mode are generally toward live-bearing or matrotrophy though this not a strictly linear progression as reversals are infrequent though plausible. We expect that improved biological sampling and phylogenetic reconstructions may yield further examples, particularly in groups displaying subgeneric transitions that are currently poorly phylogenetically resolved (e.g. within catsharks, Scyliorhinidae) or among reproductive modes which may be difficult to differentiate, such as mucoid histotrophy (Table S1).

We find no support for live-bearing being ancestral nor evidence of a high rate of reversals to egg-laying from live-bearing that this would necessitate, as suggested previously.^17^ Interestingly, a similar controversy has occurred in squamates, a group with highly labile reproductive modes. The most likely model suggested an early origin of viviparity with a high rate of reversals^18^, though this result has been questioned based on the choice of phylogenetic hypotheses and morphological features of live-bearing lineages.^1,19,20^ Consequently, support remains for the conventional hypothesis that the ancestral state was oviparity.^20^ Live-bearing chondrichthyans, particularly lecithotrophic live-bearers, develop within an egg envelope or candle which is similar to, but thinner than, that found in egg-laying species.^12,21^ This retention of the morphological machinery for egg production could make reversals to egg-laying feasible, and may allow for some of the subgeneric reproductive diversity seen within catsharks (Scyliorhinidae). But these data, along with the emerging consensus in reptiles, suggests that egg-laying is the ancestral vertebrate condition – colloquially the egg came before the chicken.

Matrotrophy appears to have evolved independently from lecithotrophic live-bearing at least 15 times with one reversal: (a-c) one to three origins within guitarfish and wedgefish (Rhinopristiformes), (d) basal to stingrays (Myliobatiformes), (e-f) one to three origins within sleeper sharks (Somniosidae), (g) great lanternsharks (*Etmopterus princeps*), (h) tawny nurse shark (*Nebrius ferrugineus*), (i) mackerel sharks (Lamniformes), (j) Pseudotriakidae, (k-m) one to three origins within houndsharks (Triakidae), and (n) base of requiem sharks (Carcharhinidae; Figure 1A). There was evidence of a single instance of reversal from matrotrophy to lecithotrophic live-bearing in the sharptooth houndshark (*Triakis megalopterus*). Overall transitions from egg-laying to live-bearing and to matrotrophy occurred at higher rates then reversals across all partitions that contained multiple reproductive modes (Figure 1C,D)

Transitions between matrotrophic modes can also occur, for example in hound sharks (Family Triakidae) muccoid histotrophy can be used in lieu of, or in concert with, placentotrophy^22^ – resulting in intergeneric reproductive diversity.^23^ Similarly, lecithotrophic live-bearing and some forms of matrotrophy, particularly muccoid histotrophy found in Rhinopristiformes and some sharks, may not represent discrete character states but rather a continuum that will be revealed only with more detailed histological work in this lineage. Muccoid histotrophy can be difficult to distinguish due to a paucity of uterine morphological specializations and accurate measurements of ash-free dry weight of embryos and ova. In several species of squaliform sharks the change in organic mass from ovum to embryo has been used to identify muccoid histotrophy^24,25^, though there is uncertainty given the exact threshold value of change in mass that should be used to distinguish between modes.^26^ Thus, large groups predominantly composed of lecithotrophic live-bearing species (i.e. Rhinopristiformes, Squalimorphii) may actually contain a greater diversity of maternal investment than currently measurable or dealt with in this analysis, and future analyses may reveal a greater number of transitions or more accurately define these currently discrete characters along a quantitative gradient of matrotrophic contribution. Despite this uncertainty, chondrichthyans still exhibit remarkably labile reproductive modes compared with other vertebrate groups, more similar to the transition rates seen in the much larger clade Squamata (∼10,000 species with >150 origins of live-bearing and 6 origins of matrotrophy).^1^

### Evolutionary covariation of reproductive mode

In sharks, live-bearing and maternal investment has covaries with larger body size, tropical latitudes, and increasing latitudinal range, while there is little evidence of covarying with depth.

#### (1) Live-bearing and matrotrophy evolve with increasing maternal size

Reproductive mode was related to body size, such that larger bodied species had a higher probability of live-bearing and matrotrophic investment (Figure 2A). We found positive covariation with body size, using the threshold model to test for evolutionary covariance between discrete values of reproductive mode and continuous ecological traits, (median = 0.16, 95% CI = 0.09 to 0.22; Effective Sample Size = 3351) indicating transitions in reproductive mode are more prevalent in lineages with larger body size (Figure 2A). The relationship with larger body size was slightly stronger for the transition from egg-laying to live-bearing (0.37 [0.17 to 0.55]; ESS = 3332) than for lecithotrophy to matrotrophy (0.3 [0.14 to 0.48]; ESS = 3600). We speculate that body size may, in effect, be capturing allometric differences in predation pressure and access to food resources, which have been hypothesized as drivers of reproductive evolution in smaller more tractable freshwater livebearers^6,27,28^. Improved foraging ability associated with increasing body size may also allow for transitions to greater degrees of maternal investment as matrotrophic forms of reproduction, particularly histotrophy and placentotrophy present greater demand for consistent energetic intake on mothers.^29,30^ We hypothesize that infrequent episodic predation and scavenging in deep sea species may preclude matrotrophy which are predominantly egg-laying or lecithotrophic live-bearing.^29^ One prediction based on food limitation would be that matrotrophic females could alter offspring size or number in response to food availability. While we see evidence of litter manipulation in chondrichthyans^31^, the opposite has been demonstrated in teleosts.^32^ It seems that, in chondrichthyans at least, the origin of live-bearing necessitates an increase in body size to accommodate retained embryos throughout gestation in limited internal body space, if larger offspring size is optimal say due to elevated predation risk in the tropics.^33^ As a result, lecithotrophic live-bearing sharks typically have fewer, but larger, pups that are presumably subject to less predation pressure and juvenile mortality.^12,29,34^

#### (2) Live-bearing and matrotrophy evolve in wide-ranging tropical species at low latitudes

Live-bearing species are more prevalent in the tropics, specifically transitions from egg-laying to live-bearing are more prevalent in lineages at lower latitudes (minimum latitude: -0.68 [-1.25 to -0.01], ESS = 2915, Figure 2C; median latitude: -0.46 [-0.9 to -0.03]; ESS = 3189; Figure 2D). However, there was little evidence that transitions in reproductive mode are related to either median latitude (-0.06 [-0.16 to 0.04], ESS = 3600) or median depth (0.06 [-0.04 to +0.15]; ESS = 3600; Figure 2 B,D). Transitions to live-bearing and in reproductive mode were more prevalent in lineages with larger latitudinal ranges (0.058 [0.08 to 1.0], ESS = 2580; 0.11 [0.04 to 0.18], ESS = 3956; Figure 2E). Despite detecting a relationship between reproductive mode and body size and latitude in extant chondrichthyans^29^, our ability to test for evolutionary covariation is hindered by the phylogenetic clustering of traits and the use of depth and latitude as proxies for environmental temperature and resource availability. As a result, it is currently difficult to disentangle evolutionary hypotheses, as large lineages with similar character states may reflect a single origin and subsequent coinheritance rather than a functional evolutionary covariation.^35^ Rays exhibit an interesting transition – consistent with the relationship we have revealed in sharks – from deep cold-water egg-laying skates (either found deep in the tropics or throughout depths in temperate and polar waters) to shallow shelf and coastal live-bearing lineages, including: electric rays (Torpediniformes), guitarfishes, wedgefishes, and sawfishes (Rhinopristiformes), and matrotrophic stingrays (Myliobatiformes). However, our power to test for plausible correlations between reproductive evolution and depth or latitude due to thermal physiology^36^, predation^37^, and productivity^29^ are limited because the batoid lineage contains a single origin of live-bearing and only one certain origin of matrotrophy. Nevertheless, tropical rays (*Himantura* spp.) paradoxically have much slower maximum population growth rates than predicted based on metabolic theory, possibly due to their large relative offspring which are thought to have evolved in response to predation risk. In turn the evolution of large offspring would only be possible through the evolution of maternal gigantism in tropical rays.^39^

### Viviparity-driven conflict versus novel ecological opportunity

Overall, the evolution of live-bearing and matrotrophy is associated with relatively greater diversification (Figure 3A-C), mainly due to a high relative extinction rate in egg-laying species (Figure 3A,B). The evolution of matrotrophy is associated with greater diversification than egg-laying (2.4 times), and there is weak evidence for greater diversification than lecithotrophic live-bearing lineages (1.27 times; Figure 4A). Egg-laying lineages have high turnover driven by high extinction rates because speciation rate is greatest in egg-laying lineages (mean 0.046 lineages/MY) compared with lecithotrophic live-bearing (mean 0.03 lineages/MY) and matrotrophic lineages (0.026 lineages/MY; Figure 3A-C).

Within the three radiations, the connection between reproductive mode and diversification is more nuanced. Within the requiem sharks (Carcharhiniformes), egg-laying lineages have a greater diversification rate than lecithotrophic live-bearing lineages (Figure 3D-F, Figure 4B), driven by high speciation in egg-laying cat sharks that are found mainly in deepwater. The “viviparity driven conflict” hypothesis predicts an increase is speciation associated with the evolution of matrotrophy, specifically placentation, as this provides a potential arena for genomic conflict between parent and offspring during pregnancy. This genomic conflict can drive sexual selection and increase the evolution of pre- and postcopulatory reproductive isolation.^5^ In poecilid fishes, increased rates of speciation are associated with pre-copulatory mechanisms, specifically male sexually selected traits, while post-copulatory mechanism (i.e. the placenta) had no effect on speciation.^3^ Likewise, we did not find support for differences in speciation associated with reproductive modes (Figure 4 A,D). Chondrichthyans do not exhibit male sexually selected traits, yet there is mounting evidence of post-copulatory mechanisms of sexual selection in sharks such as cryptic female choice and possible superfetation.^31,38^ This is a notable contrast to other fishes, and may highlight a slower rate of reproductive isolation following speciation in chondrichthyans as pre- and post-copulatory mechanisms of reproductive may not be the rate limiting factor for speciation.^14^ Reproductive isolation may be less pronounced in chondrichthyans, and indeed chondrichthyans lack secondary sexual characteristics (such as behavior, dichromatisms, and ornamentism), and there is increasing evidence of viable offspring in congeners.^40–43^ Instead novel ecological opportunities appear to be a stronger driver.

Across chondrichthyans there appears to be evidence that radiations into novel habitats rather than of reproductive mode generally affects diversification rates, such as into cold water^44^ or novel and highly dynamic fragmented freshwater habitats.^45–47^ Notably, speciation is particularly high in both the skate radiation (0.078 lineages/MY; Figure 3G) and the South American freshwater stingray radiation (0.070 lineages/MY; Figure 3J), potentially reflecting their colonization of novel polar/temperate shallows and tropical deep-water and freshwater habitats, respectively. While genomic or conflicts or sexual selection may be stronger drivers of speciation in teleosts, they are likely a weak driver of speciation compared to ecological forces across chondrichthyans.

The elevated diversification rates seen in skates (Rajiformes), South American Freshwater stingrays (Potamotrygonidae), and ground sharks (Carcharhiniformes) appear to be related to colonization of new ecological space. However, in each case, the ecological space is different – skates radiated into coldwater (polar/temperate shallows and tropical deepwater) habitats with the opening up of the Atlantic Ocean^48^, subsequent isolation of ocean basins^49^, limited egg and adult dispersal ability^48^, and a high degree of spatial niche differentiation.^50,51^ By contrast the Neotropical potamotrygonid stingrays (and other fish lineages) colonized freshwater upon the closure of the isthmus of Panama^45,46,52,53^, radiating into novel and highly fragmented freshwater habitat potentially facilitated by the emergence of unique morphological adaptations.^54^ The increased diversification in ground sharks is driven by both reinvasion of deepwater by catsharks (Scyliorhinidae) and speciation in endemic hotspots, such as South Africa, and the colonization of shallow water coastal habitats (coral reefs and associated inshore and estuarine habitats) by requiem sharks (Carcharhinidae).^44^ At present, the weight of evidence suggests that clade-specific increases in diversification rate are associated with new ecological space, rather than being systematically driven by reproductive evolution *per se* – nevertheless, we await stronger tests of this hypothesis when more extensive phylogenies become available. Future investigations of genomic conflict-driven speciation may require refined measurements of ecological variation^55^ or life history^56^ as drivers of diversification with tests restricted to smaller groups.

## Conclusions

The evolution of chondrichthyan reproductive modes, ranging from egg-laying to live-bearing and matrotrophy, appears strongly related to body size and temperature-related biogeography. While patterns of species diversification in three major radiations appear to be more strongly driven by colonization of novel habitats in contrast to teleosts, the evolution of the diversity of reproductive modes remains a fruitful area of research. Parent-offspring conflict over resources during development and subsequent antagonistic coevolution is an intriguing potential driver of reproductive mode evolution worthy of further investigation through genomic methods. In particular, the apparent slow evolution of reproductive isolation following speciation, in contrast to other fishes, hints at a different mechanism ultimately regulating the rate of speciation between shark and ray clades. Chondrichthyans are an ideal group to test for this given the diversity of reproductive modes and the frequency of polyandry, though this requires a better understanding of maternal-fetal interactions across a wider range of species.^31^ Future research could focus on improved measures of maternal investment, particularly for identifying the continuum on which lecithotrophic live-bearing and histotrophic matrotrophy may be expressed. Combined with further refinement of phylogenetic hypotheses with more extensive taxon sampling will help to clarify patterns of energetic investment, the degree of income breeding in live-bearing species, and inter- and intra-specific plasticity. More generally, we anticipate that this taxon will yield profound insights into the interplay between reproductive life history evolution, ecology, and the biogeographic patterning of species diversity across the Earth’s oceans.

## Acknowledgements

We thank A. Mooers and B. Crespi for helpful discussion regarding this study. We thank J. Hadfield and L. Revell for their guidance on the analysis of evolutionary covariation. This study was funded by the Natural Science and Environment Research Council Discovery grants to NKD and MWP, and a Canada Research Chair to NKD. KEY acknowledges funding from UNCW and a CMS Pilot Project Grant during the writing of this manuscript.

## Author Contributions

CGM, KEY and NKD conceived the study; CGM and MWP designed the analytical approach; CGM performed the analyses; CGM, MWP, KEY and NKD drafted the manuscript. All authors gave final approval for publication.

## Declaration of Interests

We have no competing interests.

## Inclusion and Diversity

We support inclusive, diverse, and equitable conduct of research.

## RESOURCE AVAILABILITY

### Lead Contact

Further information and requests for resources and reagents should be directed to and will be fulfilled by the Lead Contact, Christopher Mull.

### Materials Availability

Phylogenetic trees have been deposited at http://vertlife.org/sharktree^13^, and trait information has been deposited at http://www.sharkipedia.org^57^, and are publicly available as of the date of publication, a list of species can be found in the supplementary material.

### Data and Code Availability

All phylogenetic trees are available at www.vertlife.org/sharktree.^13^ All life history and trait information is available at www.sharkipedia.org.^57^

## EXPERIMENTAL MODEL AND SUBJECT DETAILS

### Ethics Statement

No live animals or human subjects were used in this study.

### Data Availability

All phylogenetic trees are available at www.vertlife.org/sharktree, and all trait information is available at www.sharkipedia.org.

## METHOD DETAILS

### Trait Data and Phylogeny

Data on the reproductive mode and habitat type were collected for the 610 chondrichthyan species in our phylogeny, from primary literature and species catalogues.^58–60^ Chondrichthyans exhibit eight distinct reproductive modes^2^, though we focus the evolution of live bearing and maternal investment, therefore species were categorized into three distinct modes: egg-laying, lecithotrophic live-bearing, and matrotrophic live-bearing where embryos are nourished via the initial yolk-sac investment and additional maternal contributions during gestation (oophagy, histotrophy, and placentotrophy). We note that lecithotrophic live-bearing has also been called yolk-sac viviparity, aplacental viviparity, or ovoviparity.^10^ We collected data on maximum body size and depth ranges (minimum, mean, median, and maximum) from species field guides and catalogues and primary literature. Minimum and maximum latitudinal range was collected from species geographic range maps from the International Union for the Conservation of Nature (IUCN) Red List of Threatened Species database.^58,59^ Median latitude was calculated as the midpoint between minimum and maximum latitude, and was expressed as an absolute value to represent distance north or south from the equator. All continuous trait values were standardized, centered, and divided by two standard deviations, using the rescale function in the arm (version 1.9-3)^61^ package prior to analyses to facilitate comparison of coefficients.

We conducted all analyses using a distribution of trees from a new 610 species chondrichthyan molecular phylogeny.^13^ This phylogeny covers 51% of all known species from every order, 98% of families, 88% of genera, and all described character states are represented. Because the distribution of trees represents a gradient of variation in root and node dating for the maximum likelihood tree, we sequentially selected 21 trees, every 25^th^ tree from one to 500 to account for the full range of temporal calibrations. Results were pooled across all trees.

### Ancestral State Reconstruction and Diversification

We reconstructed the evolutionary origins and sequence of reproductive mode and habitat while estimating state dependent diversification rates using the multistate speciation and extinction (MuSSE) method with maximum likelihood implemented with musse^62^ in the diversitree (version 0.9-9) package in R.^63^ Rabosky and Goldberg ^64^ pointed out that state dependent diversification models (including MuSSE) are susceptible to inflated false positives when there is unmodeled heterogeneity in diversification rates – thereby associations can be falsely detected between rates and states even when the diversification dynamics are unrelated to the traits being considered. To minimize false positives, we adopted the analytical approach of Uyeda et al.^65^ and integrated a hypothesis-testing approach (i.e., MuSSE) with a more data-driven approach (what Uyeda et al. refer to as “phylogenetic natural history”); in this way, we are able to tease out the signal for our hypotheses that our focal traits are influencing diversification rates while conditioning on the fact that there is likely substantial background variation in these rates across the tree.

More specifically, we first used the medusa algorithm (as implemented in Geiger v2.0.6^66^ to detect background variation in diversification rates unrelated to our traits of interest. First, we infilled those species missing from the molecular phylogeny (n = 582) using taxonomic constraints so that tips represented unresolved clades. Importantly for earlier divergences, our sampling did not miss any variability in reproductive mode (e.g. there are no live-bearing chimaeras) so missing samples will not affect our conclusions. Under some circumstances, medusa may not reliably correctly identify the placement of shifts.^67^ However, this issue does not pertain to our analysis because we are not making any inferences about specific events or specific clades and are only interested in detecting broad-scale differences in diversification dynamics. Our Medusa analysis only revealed three clades with consistent increases in diversification rates across all trees: skates (Rajiformes), South American freshwater stingrays (Potamotrygonidae), and ground sharks (Carcharhiniformes). For the MuSSE analyses, we then partitioned the tree into four diversification regimes (these three known radiations plus the background) and fit a MuSSE model in which the diversification rate parameters were allowed to vary among these four partitions. We also assigned each partition its own sampling fraction based on the most up-to-date taxonomic treatment.^13,68^ Ideally, we would have fit an integrated model that included heterogeneity in rates due to both the trait and to unmodeled “background” variation^65,69^ but unfortunately no such approach is available for multi-state traits.

As we described above, speciation (λ), extinction (μ), and transition rates (*q*) were estimated for each of the four partitions separately. Because the estimations of extinction rates from molecular phylogenies can be difficult^70^, we ran two models: with state dependent extinction rates (1) unconstrained (μ_egg-laying_ ≠ μ_live-bearing_ ≠ μ_matrotrophic_) and (2) constrained to be equal (μ_egg-laying_ = μ_live-bearing_ = μ_matrotrophic_). We report findings from the unconstrained model as there was no significant difference between models, and we are interested in how variation in this rate may affect overall diversification. Additionally, the main difficulty with estimating extinction rates arises from unaccounted for diversification rate heterogeneity, which we have minimized by *a priori* identification with medusa and subsequent partitioning. We report speciation (λ), extinction (μ), net diversification (*r* = λ – μ), and transition rates (*q*). We treated reproductive mode as an ordinal multistate character and accordingly did not allow unlikely transitions such as directly between egg-laying and matrotrophic live-bearing. Models were run for 10,000 generations with the first 1,000 generations discarded as burn-in, using an exponential prior with a rate of 1/(2*r*) where *r* is the character state independent diversification rate.^63^ We checked that all parameter estimates had effective sample sizes (ESS: the number of independent draws from an MCMC chain) greater than 200. To account for autocorrelation in the estimates of state dependent diversification, we also examined the posterior distribution of differences in state dependent rates across all chains.

### Evolutionary Covariation with Ecological Traits

We used a threshold model to test for the evolutionary covariation between reproductive mode and continuous ecological traits. The threshold model assumes that state changes in an ordinal variable (e.g. egg-laying to live-bearing to matrotrophy) occur when a threshold value of an underlying continuous latent variable, such as body size, is reached. Thus, it can be used to model the evolutionary covariation between ordinal and continuous traits.^71^ Accurate estimation of evolutionary covariation requires a suitable number of transitions and distribution of traits across the phylogeny.^35^ We focus on sharks (superorders Galeomorphii and Squalomorphii; n = 292) to evaluate evolutionary covariance between reproductive transitions and three ecological traits (body size, depth, and latitude), because there is only one transition in parity and few appearances of matrotrophy within Chimaeriformes and rays (Batoideii). We estimated the evolutionary covariance using Bayesian methods, sampling from the posterior distribution using a special Reduced Animal Model implemented in a mixed effects modeling framework while accounting for phylogeny, using the package MCMCglmmRAM (version 2.24) in R.^72^ This approach is equivalent to estimating evolutionary covariance using the threshold model.^71,73^ These models are a special case of generalized linear mixed effects models where heritability, akin to Pagel’s λ, is set to a value of one corresponding to Brownian motion with respect to the phylogenetic tree.^74,75^ Twenty chains were run for 2 million generations with the first 200,000 iterations discarded as burn-in, using priors with an inverse-Wishart distribution and the residual covariance matrix set to zero.^72^ Samples were drawn every 500 iterations to avoid temporal autocorrelation in parameter estimates. Chains were visually inspected to ensure convergence using coda (version 0.19-4)^76^, and posterior samples were summarized to generate mean and 95% highest posterior densities (HPD) with effective samples sizes greater than 1000. Models were run using three different treatments of reproductive mode with the threshold family: binary parity mode (egg-laying versus live-bearing), binary embryo trophic mode (lecithotrophic versus matrotrophic), and ordinal multi-state reproductive mode (egg-laying, lecithotrophic live-bearing, and matrotrophic live-bearing).

## Notes

http://vertlife.org/sharktree/

http://www.sharkipedia.org

### Competing Interest Statement

The authors have declared no competing interest.

### Summary of Updates

This manuscript has been reformatted and new citations have been added. Additionally we provide further discussion of the rate limiting mechanisms of diversification and reproductive isolation across fishes.

